# Biodiversity and Antimicrobial Potential of Bacterial Endophytes from Halophyte *Salicornia brachiata*

**DOI:** 10.1101/2020.07.15.203612

**Authors:** Sanju Singh, Vishal Ghadge, Pankaj Kumar, Doniya Elze Mathew, Asmita Dhimmar, Harshal Sahastrabudhe, Yedukondalu Nalli, Mina R. Rathod, Pramod B. Shinde

## Abstract

Untapped natural habitats like halophytes, marsh land, and marine environment are suitable arena for chemical ecology between plants and microbes having environmental impact. Endophytes constitute an ecofriendly option for the promotion of plant growth and to serve as sustainable resources of novel bioactive natural products. The present study focusing on biodiversity of bacterial endophytes from *Salicornia brachiata*, led to isolation of around 350 bacterial endophytes. Phylogenetic analysis of 63 endophytes revealed 13 genera with 29 different species, belonging to 3 major groups: Firmicutes, Proteobacteria and Actinobacteria. 30% isolates belonging to various genera demonstrated broad-spectrum antibacterial and antifungal activities against a panel of human, plant, and aquatic infectious agents. An endophytic isolate *Bacillus amyloliquefaciens* 5NPA-1, exhibited strong in-vitro antibacterial activity against human pathogen *S. aureus* and phytopathogen *X. campestris*. Investigation through LC-MS/MS-based molecular networking and bioactivity-guided purification led to the identification of three bioactive compounds belonging to lipopeptide class on the basis of ^1^H- and ^13^C-NMR and MS analysis. To our knowledge, this is the first report studying bacterial endophytic biodiversity of *Salicornia brachiata* and isolation of bioactive compounds from its endophyte. Overall, the present study provides insights into the diversity of endophytes associated with the plants from the extreme environment as rich source of metabolites with remarkable agricultural applications and therapeutic properties.

## INTRODUCTION

Ecological interactions are responsible for providing prime ecosystem services. Plant mediated interactions and structure of natural communities go hand in hand as they potentially link organisms of different trophic levels and add chemical complexity within communities leading to surging catalog of compounds. Endophytes, impart protection to plants against various abiotic or biotic stress tolerance by production of plant hormones, bioavailability of nutrients, and antagonistic action to phytopathogens in turn, sustaining on the nutrients provided by plants thereof [1]. Endophytic bacteria facilitates plant growth and developments through various processes including nitrogen fixation, phosphate solubilization, production of hormones and siderophores, and decreasing ethylene concentration [2,3]. Along with host plant-growth cycle, endophytes also improvise their survival mechanisms during their continuous efforts to live in the host tissues [4]. Biological control within host is mediated by endophytes that promotes plant growth by protecting against the attack of phytopathogens, facilitated by the production of siderophores, antibiotics or bacteriocins [5]. It has been proved that bioactive compounds derived from plants are mostly secondary metabolic products of microbes inhabiting inside the plants symbiotically defined as endophytes [6]. Predominantly, actinobacterial and bacterial endophytes contribute heavily in plant growth promotion and agriculture management strategies via production of metabolites such as aromatic compounds, lipopeptides, plant hormones, polysaccharides, and several enzymes linked to phenylpropanoid metabolism [7].

Halophytes are plants that thrive in saline environment with salinity upto 200 mM concentration [8]. Endophytic bacteria and fungi isolated from halophytes aid hosts by altering plant hormone status and uptake of nutrient elements and/or modulating the production of reactive oxygen species through different mechanisms [9,10]. Strains of *Bacillus amyloliquefaciens* are reported to enhance plant growth promotion and also provide defense benefits against phytopathogens like *Phytophthora parasitica var. nicotiana, Fusarium oxysporum* sp., *F. graminearum, F. solani, Alternaria alternate* etc. [11,12]. Beneficial metabolites produced due to ecological interactions between endophyte-plant can be harnessed and employed for multifaceted applications in arena such as agriculture, medicine, bioremediation, and biodegradation. Despite their beneficial characters, the research regarding endophytes from plants inhabiting extreme environments is still at an early stage with respect to diversity of endophytes, their functional roles, and bioprospecting for bioactive compounds.

*Salicornia brachiata* is a halophyte with medicinal properties having salty marsh lands as the natural habitat getting exposed to extremities of salinity, heat, temperature, and humidity [13]. *S. brachiata* was reported to harbor plant-growth promoting microorganisms [14,15]. It can be hypothesized that *S. brachiata* being a plant inhabiting in extreme environment, must have evolved ways to harbor diverse kinds of endophytes apart from plant-growth promoters and to employ those endophytes for protection from herbivores and pests including phytopathogenic fungi and insects, competing for nutrients including trace elements. Motivated by this hypothesis, present study investigated diversity of bacterial endophytes associated with the halophyte *S. brachiata* sampled at three distinct locations at Gujarat coast, India. Further, their bioactive potential was also studied against a panel of human, plant, and aquatic infectious agents. Repeated chromatographic separation resulted in isolation of three compounds from the endophyte *B. amyloliquefaciens*. This may be the first comprehensive study evaluating bacterial endophytic biodiversity of the halophyte *Salicornia brachiata*.

## METHODS

### Collection of plant material and isolation of endophytes

Healthy and fresh plant samples of *S. brachiata* were collected randomly from three different sites i.e. New port (N 21° 45’ 15.7”, E 072° 14’ 01.4’’), Sartanpar port (N 21° 17’ 52.1’’, E 072° 66’ 25.5”), Victor port (N 20° 58’ 53.2”, E 071° 33’ 21.2’’), located along Gujarat coast, India (**Figure 1**). Samples were placed in sample bags and stored on icebox right after sampling to preserve microbial flora and then the plant material was transported to laboratory and processed immediately.

**Fig. 1.**
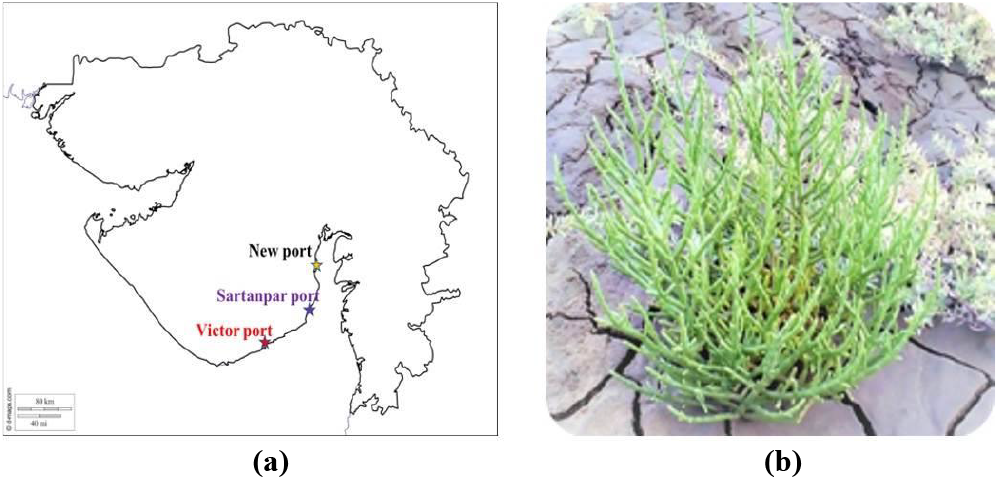
Sampling location. (a) Location Map of Gujarat coast, India showing three sampling sites 1) New port, 2) Sartanpar port, 3) Victor port. (b) Morphology of *Salicornia brachiata* plant in natural marsh habitat

The plant material was surface sterilized followed by different treatments to enhance the probability of maximum number of novel bacterial or actinobacterial species. Surface sterilization of the samples was carried out by reported method [16]. Sterilized samples were aseptically fragmented into small pieces and directly placed on eight selective media prepared with cycloheximide and nystatin at concentration of 50 mg/ml to inhibit fungal growth. Sterilized samples were given six different pre-treatments (**Table I**) and plates were incubated at 30 °C for 2 to 8 weeks. Control plates inoculated with last wash of sterilization procedure was incubated to check effectiveness of surface sterilization in triplicate. Periodic growth analysis and subsequent sub-culturing for purification of isolates was performed. Glycerol stocks of purified strains were prepared and stored at −80 °C.

**Table I.** Location, Pre-treatments and Media used in the study.

| Sites of sampling | Pre-treatments | Media |
| --- | --- | --- |
| 1. **New port** | 1. Sample grinded by mortar pestle, diluted and spread. | 1. Nutrient Agar (NA) |
| 2. **Sartanpar port** |  | 2. Zobell Marine agar (ZMA) |
| 3. **Victor port** | 2. Moist incubation of grinded sample with CaCO<sub>3</sub> for 7-14 days and then spread. | 3. Tryptic Soya agar (TSA) |
|  | 3. Rehydration and centrifugation method. | 4. Starch Casein Agar (SCA) |
|  | 4. Dry heating the plant at 80-100 °C. | 5. Yeast Malt Agar (ISP 2) |
|  | 5. Treatment with 1.5% (w/v) phenol. | 6. Actinomycetes isolation agar (AIA) |
|  | 6. Treatment with 0.01% potassium dichromate. | 7. Inorganic Salts Starch agar 4 (ISP 4) |
|  |  | 8. Tap water Yeast Extract Agar (TWYA) |

### Identification of endophytes

DNA isolation was performed using reported protocol with some modification [17]. Briefly, 2 ml of 24 h grown bacterial culture (48 h for slow growing bacteria) was centrifuged for 2 min at 13000 rpm. Cell pellet was resuspended in 600 μl TE buffer (10 mM Tris base, 1 mM EDTA, pH 8.0). 100 μl of 10 mg/ml lysozyme was added and incubated at 37 °C for 1 h followed by addition of 20 μl of 20 mg/ml Proteinase K and incubation at room temperature for further 1 h. After incubation, 200 μl of 10% SDS was added and kept at 55 °C for 1 h followed by purification through extraction of aqueous phase with phenol: chloroform: isoamyl alcohol (24:24:1). DNA was precipitated using 3M sodium acetate and chilled isopropanol and the obtained DNA pellet was washed by ethanol. 16S rRNA amplification was done using universal primer sequences of FD_1_ (5’–GAGTTTGATCCTGGCTCA-3’) and RP2 (5’–ACGGCTAACTTGTTACGACT-3’). Reaction mixture, consisting of DNA template – 1 μl (50 ng/μl), Primers – 0.5 μl of 10 μM, dNTP – 0.2 mM and Taq polymerase – 1.25 units, was prepared and the reaction was performed in Thermocycler (Bio-Rad T100) with conditions: Initial denaturation – 95 °C for 5 min, 34 cycles of 95 °C for 30 s, 58 °C for 30 s and 55 °C for 1 min and final amplification for 5 min.

Amplification of 16S rRNA gene was confirmed by gel electrophoresis; subsequently PCR products were purified with Qiagen PCR purification kit and sequenced by Macrogen Inc. Korea. The obtained sequences were trimmed to align in BioEdit software (version 7.0.5.3) [18] and consensus sequence created was searched in NCBI GenBank database using BLAST. Further, 16S rRNA sequences were aligned and used to construct maximum likelihood phylogenetic tree using molecular evolutionary genetic analysis MEGA 6 software [19].

### Statistical data analysis

Simpson index of diversity and Shannon-Wiener’s diversity index were calculated to determine endophytic diversity obtained from samples of all the three locations [20].

Simpson’s index of diversity (D) gives the probability that two individuals selected at random will belong to the same species and was calculated using formula as given in the following equation:

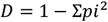

Shannon-Wiener diversity index (H) determines actual diversity of the bacterial endophytes, and was calculated using the following equation:

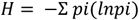

where p_i_ = n/N;
n = total number of organisms of species ‘i’.

N = total number of organisms of all species.

Shannon evenness (E) was calculated as H′/H_max_, where H_max_= ln(S), with S as the total number of species in the sample. Further, to determine qualitative wealth species richness was also calculated by S/√N.

### Antimicrobial potential of endophytes

A panel of 8 reference human, plant, and marine pathogens comprising of *Staphylococcus aureus* MCC 2043, *Bacillus subtilis* MCC 2049, *Mycobacterium smegmatis* MTCC 6, *Escherichia coli* MCC 2412, *Candida albicans* MCC 1152, *Xanthomonas campestris* NCIM 5028, *Fusarium oxysporum* NCIM 1008, *Alteromonas macleodii* NCIM 2815 was employed to determine bioactive potential of all the endophytic isolates. 10 ml culture of endophytes in 50 ml tubes was done for a period of 14 days at 30 °C, 180 rpm shaking condition, in-between assessing it at an interval of 7 days for the antimicrobial activity. Whole cell culture and supernatant (obtained after centrifugation) were used to perform well diffusion assay [21] for determination of bioactivity.

### Preparation of crude extracts

In order to confirm endophytes play important role in survival of halophyte *S. brachiata* in extreme climate by protecting against pathogens, crude extracts of isolates which were found bioactive in primary screening were prepared. Briefly, 250 ml culture broth in 1000 ml Erlenmeyer flask in particular medium and incubation conditions specific for an isolated strain was done. After incubation period, culture broths were extracted twice with equal volume of ethyl acetate. Organic phase was collected over anhydrous sodium sulfate and concentrated under rotary evaporator to yield dark-brown colored crude extracts. The activity of crude extracts of isolates bioactive against *S. aureus* MCC 2043 and *X. campestris* NCIM was confirmed through disk diffusion assay by loading 1 mg of extract on the disk following Clinical and Laboratory Standards Institute (CLSI) protocols [22].

### LC-MS/MS based molecular networking

LC-MS/MS data of the crude extract of *B. amyloliquefaciens* 5NPA-1 was studied with GNPS (Global natural products social molecular networking) (https://gnps.ucsd.edu) [23]. Raw data received from Agilent 6545 Q-TOF LC/MS was converted into GNPS compatible mzXML format using MSconvert application (version 3.0.19317-0ef6e44d0) in Proteowizard suit [24]. The converted file was uploaded to GNPS server (massive.ucsd.edu) using WinSCP FTP client. Molecular networking was run as classical molecular networking work flow (METABOLOMICS-SNETS-V2) with activated MS-Cluster. Parameters for input algorithm were set as: precursor ion mass tolerance 2.0 Da, fragment ion mass tolerance 0.5 Da, minimum cosine score for an edge formation by two consensus MS/MS spectra as 0.7, minimum number of common fragment ions as 6 (no. of fragment shared by two separate consensus MS/MS spectra in order to be connected by an edge in the molecular network), minimum cluster size was 2, and edges between two nodes were considered only if both nodes were within top 10 most similar nodes of each other. Molecular network file was downloaded in GraphML format and visualized using Cytoscape (version 3.7.2) [25].

### TLC bioautography

Ethyl acetate extract of endophyte *B. amyloliquefaciens* 5NPA-1 was subjected to thin layer chromatography (TLC) analysis over analytical aluminium silica gel 60 TLC plate for separation of metabolites to obtain R_f_ value of active fraction. The crude extract was dissolved in methanol to get a concentration of 10 mg ml^−1^ and 1 mg was loaded on TLC plate which was developed with solvent system comprising of methanol and chloroform in the ratio 12:88. Separate bands were observed under short wavelength UV light and their R_f_ values were calculated. To perform bioautography, 1.5 ml of 1% mueller hinton agar (MHA) was spread on the developed TLC plate of size 10 × 2 cm under sterile environment of biosafety cabinet and 50 μl of log phase culture of *S. aureus* MCC 2043 was spread with the help of sterile spreader. After incubation for 24 h at 30 °C, the plate was visualized by spraying iodonitrotetrazolium chloride solution (INT, 2 mg ml^−1^) pink color on plates signified cell growth whereas bands with clear zone indicated inhibition of cell growth. R_f_ values of the bands with clear zone were recorded [26].

### Isolation and identification of bioactive molecules

Purification of compounds was carried out by separating 50 mg of crude extract on preparative TLC plates (Kieselgel 60 F_254_, 25 mm, Merck) using the same solvent system as above. Based on TLC bioautography, the active portion was selectively scrapped and dissolved in methanol. The supernatant was concentrated on rotary evaporator to obtain 15 mg of yellow, viscous oil (5-PTLC2), which was analyzed by HPLC. Further, ^1^H-NMR spectrum (Bruker, Avance II 500 MHz in CD_3_OD) of 5-PTLC2 was acquired in order to identify the class of compounds. Additionally, to purify active compounds, the crude extract (2 g) was fractionated by MPLC (C18 RP-silica gel) eluting with aqueous MeOH (30% to 100%) resulting in the collection of 18 fractions (fr. 1–fr. 18) which were evaluated for antimicrobial activity against *S. aureus* MCC 2043. Fractions 11 and 13 containing bioactive compounds were further chromatographed on semi-preparative HPLC (Dionex Ultimate 3000, Thermo Scientific) using gradient mixtures of acetonitrile–water (4:1 to 9:1) on YMC column (YMC-Triart C18, 5 μm, 250 × 10 mm I.D.) to yield three compounds (**1**–**3**). The three compounds were further characterized by MALDI-MS and MALDI-MS/MS (Applied Biosystems 4800 MALDI TOF–TOF analyzer) and NMR (JEOL 600 MHz).

## RESULTS

### Collection of plant material and isolation of endophytes

*S. brachiata* samples collected from three different sites along Gujarat coast (**Fig. 1**) were processed using eight different media with six variable pretreatments resulting in isolation of 336 endophytes differentiated on the basis of morphological characters (**Fig. S1**). The high number of isolates obtained were also found to be equally diversified. Also, effectiveness of surface sterilization was established as no growth was observed on the negative control plate even when incubated for a month long period. Maximum number of bacterial isolates were obtained on medium supporting fast growing microbes i.e. 81 isolates collectively from NA, ZMA, and TSA; 52 from SCA; 52 from ISP2; 50 from AIA; 70 from ISP4; 31 from TWYA (**Fig. S2**). The diverse group of isolated actinobacteria preferred ISP4 media for their growth and pigment production.

### Identification of endophytes

Out of 336 isolates, 63 isolates obtained from different media were identified on the basis of molecular phylogeny by 16S rRNA sequencing. Evolutionary relationships of different endophytes obtained from *S. brachiata* were inferred from maximum likelihood phylogenetic tree (**Fig. 2**). On the basis of 16S rRNA sequencing results, the most predominant as well as diverse genera identified was *Bacillus* being 76% of the isolates with 15 different species. The remaining percentage accounted for various genera of actinomycetes or bacteria like *Isoptericola, Paenibacillus, Nocardiopsis, Rhodococcus, Salinicola, Jonesia, Nitratireductor*, *Enterococcus, Streptomyces, Micromonospora, Pseudomonas* and *Marinilactibacillus* (**Table II**).

**Fig. 2.**
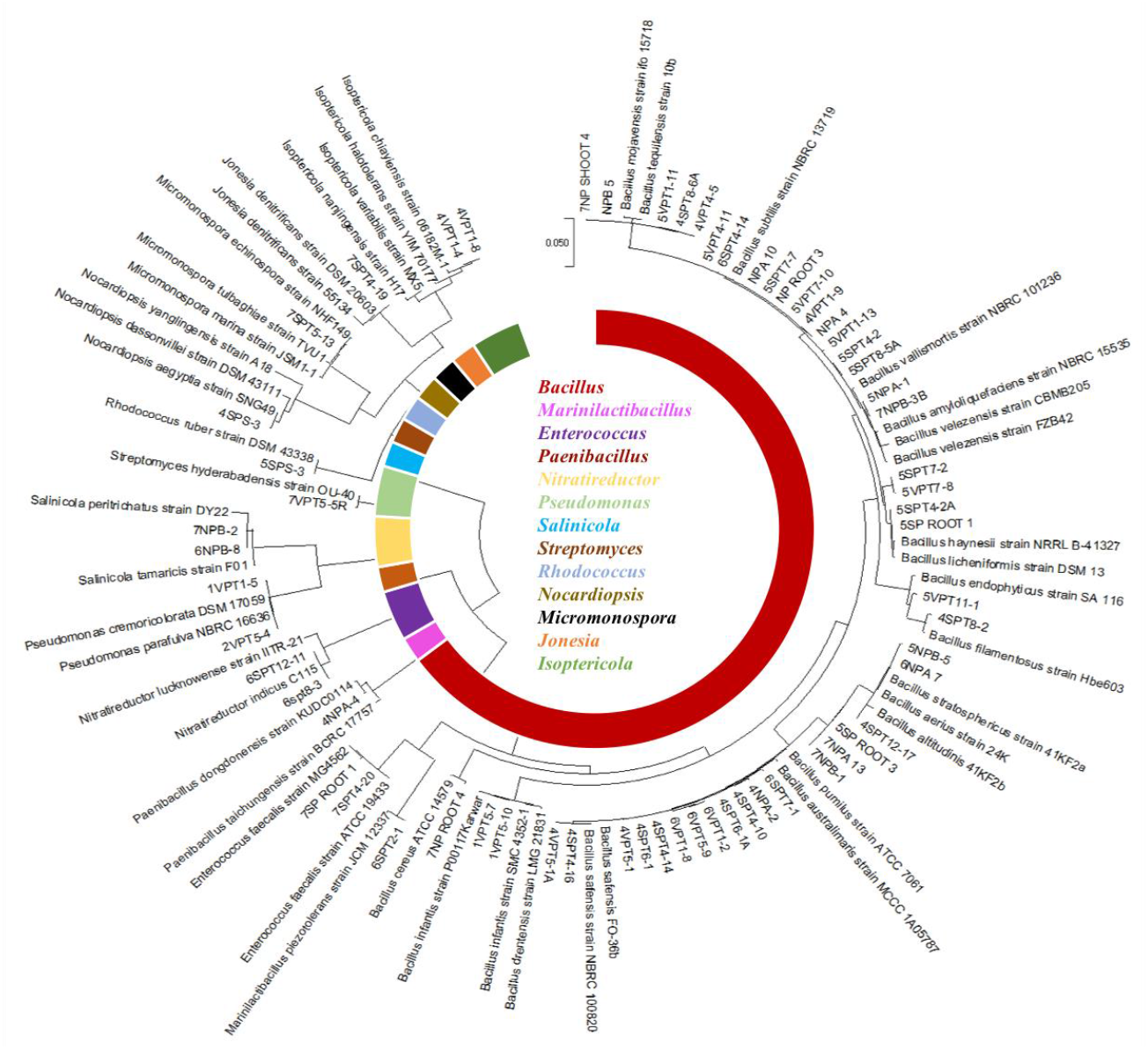
Identification of endophytes by 16S rRNA sequencing. Phylogenetic tree of 63 bioactive isolates obtained by Maximum likelihood analysis constructed using Mega 6 software depicting their phylogeny with related genera as well as pie graph displaying percentage distribution of genera obtained

**Table II.** Identification and similarity values of 16S rDNA sequences retrieved from the endophytic bacteria from *S. brachiata*.

| S.no. | Isolates | Identification | Accession number | Closest bacteria in database | Similarities (%) |
| --- | --- | --- | --- | --- | --- |
| 1 | NPROOT-3 | *Bacillus subtilis* | MT459325 | *B. subtilis* NR_027552 | 99.86 |
| 2 | NPB-5 | *B. subtilis* | MT645716 | *B. subtilis* NR_027552 | 99.93 |
| 3 | NPA-10 | *B. tequilensis* | MT645715 | *B. tequilensis* NR_104919 | 99.86 |
| 4 | NPA-4 | *B. subtilis* | MT645714 | *B. subtilis* NR_027552 | 99.93 |
| 5 | 1VPT1-5 | *Pseudomonas parafulva* | MT645710 | *Pseudomonas parafulva* NR_104280 | 99.86 |
| 6 | 1VPT5-7 | *B. infantis* | MT645711 | *B. infantis* NR_043267 | 99.63 |
| 7 | 1VPT5-10 | *B. infantis* | MT645712 | *B. infantis* NR_043267 | 99.63 |
| 8 | 2VPT5-4 | *Pseudomonas cremoricolorata* | MT645713 | *Pseudomonas cremoricolorata* NR113855 | 100 |
| 9 | 4NPA-2 | *B. safensis* | MT645719 | *B. safensis* NR_041794 | 99.79 |
| 10 | 4NPA-4 | *Paenibacillus taichungensis* | MT645718 | *Paenibacillus taichungensis* MH553940 | 99.29 |
| 11 | 4SPS-3 | *Nocardiopsis aegyptia* | MT645720 | *Nocardiopsis aegyptia* NR025889 | 99.71 |
| 12 | 4SPT4-10 | *B. safensis* | MT645721 | *B. safensis* NR_041794 | 99.78 |
| 13 | 4SPT4-14 | *B. safensis* | MT645722 | *B. safensis* NR_041794 | 99.93 |
| 14 | 4SPT4-16 | *B. safensis* | MT645723 | *B. safensis* NR_041794 | 99.79 |
| 15 | 4SPT6-1 | *B. safensis* | MT645724 | *B. safensis* NR_041794 | 100% |
| 16 | 4SPT6-1A | *B. zhangzouensis* | MT645725 | *B. safensis* NR_041794 | 99.79 |
| 17 | 4SPT8-2 | *B. filamentosus* | MT645726 | *B. filamentosus* NR_134701 | 99.64 |
| 18 | 4SPT8-6A | *B. subtilis* | MT645727 | *B. subtilis* NR_113265 | 99.93 |
| 19 | 4SPT12-17 | *B. aerius* | MT645728 | *B. aerius* NR_118439 | 99.86 |
| 20 | 4VPT1-4 | *Isoptericola chiayiensis* | MT645729 | *Isoptericola chiayiensis* NR_116696 | 99.05 |
| 21 | 4VPT1-8 | *Isoptericola chiayiensis* | MT645730 | *Isoptericola chiayiensis* NR_116696 | 99.20 |
| 22 | 4VPT1-9 | *B. subtilis* | MT645731 | *B. subtilis* NR_027552 | 99.35 |
| 23 | 4VPT4-5 | *B. tequilensis* | MT645732 | *B. tequilensis* NR_104919 | 99.86 |
| 24 | 4VPT5-1 | *B. pumilus* | MT645733 | *B. pumilus* NR_043242 | 99.71 |
| 25 | 4VPT5-1A | *B. zhangzhouensis* | MT645717 | *B. zhangzhouensis* NR_148786 | 99.35 |
| 26 | 5NPA-1 | *B. amyloliquefaciens* | MT459305 | *B. amyloliquefaciens* NR_041455 | 98.91 |
| 27 | 5SPRoot-1 | *B. paralicheniformis* | MT645735 | *B. paralicheniformis* MK517555 | 99.93 |
| 28 | 5NPB-5 | *B. aerius* | MT645734 | *B. aerius* NR_118439 | 99.78 |
| 29 | 5SPS-3 | *Rhodococcus ruber* | MT645737 | *Rhodococcus ruber* NR_118602 | 99.93 |
| 30 | 5SPT4-2 | *B. tequilensis* | MT645738 | *B. tequilensis* NR_104919 | 99.71 |
| 31 | 5SPT7-2 | *B. paralicheniformis* | MT645740 | *B. paralicheniformis* MF_321822 | 99.93 |
| 32 | 5SPROOT 3 | *B. stratosphericus* | MT645736 | *B. stratosphericus* NR_118441 | 99.93 |
| 33 | 5SPT4-2A | *B. paralicheniformis* | MT645739 | *B. paralicheniformis* MK517555 | 99.86 |
| 34 | 5SPT7-7 | *B. tequilensis* | MT645741 | *B. tequilensis* NR_104919 | 99.86 |
| 35 | 5SPT8-5A | *B. tequilensis* | MT645742 | *B. tequilensis* NR_104919 | 99.86 |
| 36 | 5VPT7-8 | *B. paralicheniformis* | MT645746 | *B. paralicheniformis* MF321822 | 99.93 |
| 37 | 5VPT11-1 | *Bacillus endophyticus* | MT645748 | *Bacillus endophyticus* KY194734 | 98.8% |
| 38 | 5VPT1-11 | *B. tequilensis* | MT645744 | *B. tequilensis* NR_104919 | 99.86 |
| 39 | 5VPT4-11 | *B. subtilis* | MT645745 | *B. subtilis* NR_113265 | 99.86 |
| 40 | 5VPT7-10 | *B. subtilis* | MT645747 | *B. subtilis* NR_112629 | 99.93 |
| 41 | 5VPT1-13 | *B. subtilis* | MT645743 | *B. subtilis* NR_113265 | 99.72 |
| 42 | 6SPT2-1 | *Marinilactibacillus piezotolerans* | MT645751 | *Marinilactibacillus piezotolerans* NR_112661 | 99.56 |
| 43 | 6NPB-8 | *Salinicola tamaricis* | MT645750 | *Salinicola tamaricis* NR_157001 | 99.34 |
| 44 | 6SPT4-14 | *B. subtilis* | MT645755 | *B. subtilis* NR_113265 | 99.79 |
| 45 | 6SPT7-1 | *B. safensis* | MT645752 | *B. safensis* NR_041794 | 99.65 |
| 46 | 6SPT12-11 | *Nitratireductor indicus* | MT645754 | *Nitratireductor indicus* NR_117518 | 99.55 |
| 47 | 6VPT1-2 | *B. safensis* | MT645756 | *B. safensis* NR_041794 | 99.79 |
| 48 | 6VPT5-9 | *B. zhangzouensis* | MT645758 | *B. zhangzouensis* NR_148786 | 99.50 |
| 49 | 6NPA-7 | *B. aerius* | MT645749 | *B. aerius* NR_118439 | 99.64 |
| 50 | 6SPT8-3 | *Nitratireductor indicus* | MT645753 | *Nitratireductor indicus* NR_117518 | 99.70 |
| 51 | 6VPT1-8 | *B. pumilus* | MT645757 | *B. safensis* NR_112637 | 99.57 |
| 52 | 7NPA-13 | *B. stratosphericus* | MT645759 | *B. stratosphericus* NR_118441 | 99.93 |
| 53 | 7NPB-1 | *B. stratosphericus* | MT645760 | *B. stratosphericus* NR_042336 | 99.86 |
| 54 | 7NPB-2 | *Salinicola tamaricis* | MT645761 | *Salinicola tamaricis* NR_157001 | 98.80 |
| 55 | 7VPT5-5R | *Streptomyces hyderabadensis* | MT645769 | *Streptomyces hyderabadensis* NR_116934 | 99.55 |
| 56 | 7NPSHOOT-4 | *B. subtilis* | MT645765 | *B. subtilis* NR_112629 | 100 |
| 57 | 7NPB-3B | *B. velezensis* | MT645763 | *B. velezensis* NR_116240 | 99.78 |
| 58 | 7SPROOT-1 | *Enterococcus faecalis* | MT645764 | *E. faecalis* NR_115765 | 99.80 |
| 59 | 7SPT4-20 | *Enterococcus faecalis* | MT645767 | *E. faecalis* NR_115765 | 99.93 |
| 60 | 7SPT4-19 | *Jonesia denitrificans* | MT645766 | *Jonesia denitrificans* NR_119162 | 99.78 |
| 61 | 7NPROOT-4 | *B. cereus* | MT645762 | *B. cereus* NR_115526 | 99.51 |
| 62 | 7SPT5-13 | *Micromonospora echinospora* | MT645768 | *Micromonospora echinospora* NR_118843 | 99.41 |
| 63 | 4NPBL | *B. subtilis* | MT447880 | *B. subtilis* NR_112116 | 99.86 |

### Statistical data analysis

α-diversity indexes, i.e. Shannon-Wiener diversity index and Simpson’s index of diversity and their components; richness and evenness were used to determine diversity of the endophytic community isolated from *S. brachiata* sampled from three different locations. Shannon-Wiener diversity index (H) was calculated as 3.028, indicating high diversity within endophytes. Species richness indicating abundance of the species in a sample was found to be 3.615, viz., greater the value higher the richness. Shannon evenness measures relative abundance of different species contributing to richness of a sample, was calculated as 0.899 signifying an even community structure [20]. In agreement with aforementioned data, Simpson’s index of diversity (D) of 0.931, demonstrate a high diversity of endophytes harbored by the host halophyte (**Table III**).

**Table III.** Diversity indices of the 63 endophytes isolated from *S. brachiata*.

| Diversity indices | Values calculated | Range |
| --- | --- | --- |
| Shannon-Wiener diversity index | 3.028 | 0 onwards |
| Simpson's index of diversity | 0.931 | 0-1 |
| Species richness | 3.615 | 0 onwards |
| Shannon evenness | 0.899 | 0-1 |

### Antimicrobial potential of endophytes

From the primary screening study, a total of 101 isolates were found to exhibit broad-spectrum antimicrobial activity against one or the other pathogen from panel of pathogens tested. 8 isolates were found to be active against *M. smegmatis* MTCC 6; 20 against *S. aureus* MCC 2043; 51 against *X. campestris* NCIM 5028; 42 against *A. macleodii* NCIM 2815; 12 against *C. albicans* MCC 1152; 14 against *F. oxysporum* NCIM 1008; 19 against *B. subtilis* MCC 2049; 5 against *E. coli* MCC 2412 (**Fig 3a**). It was observed that some endophytes displayed inhibition activity against two or more pathogens (**Table IV**).

**Fig. 3.**
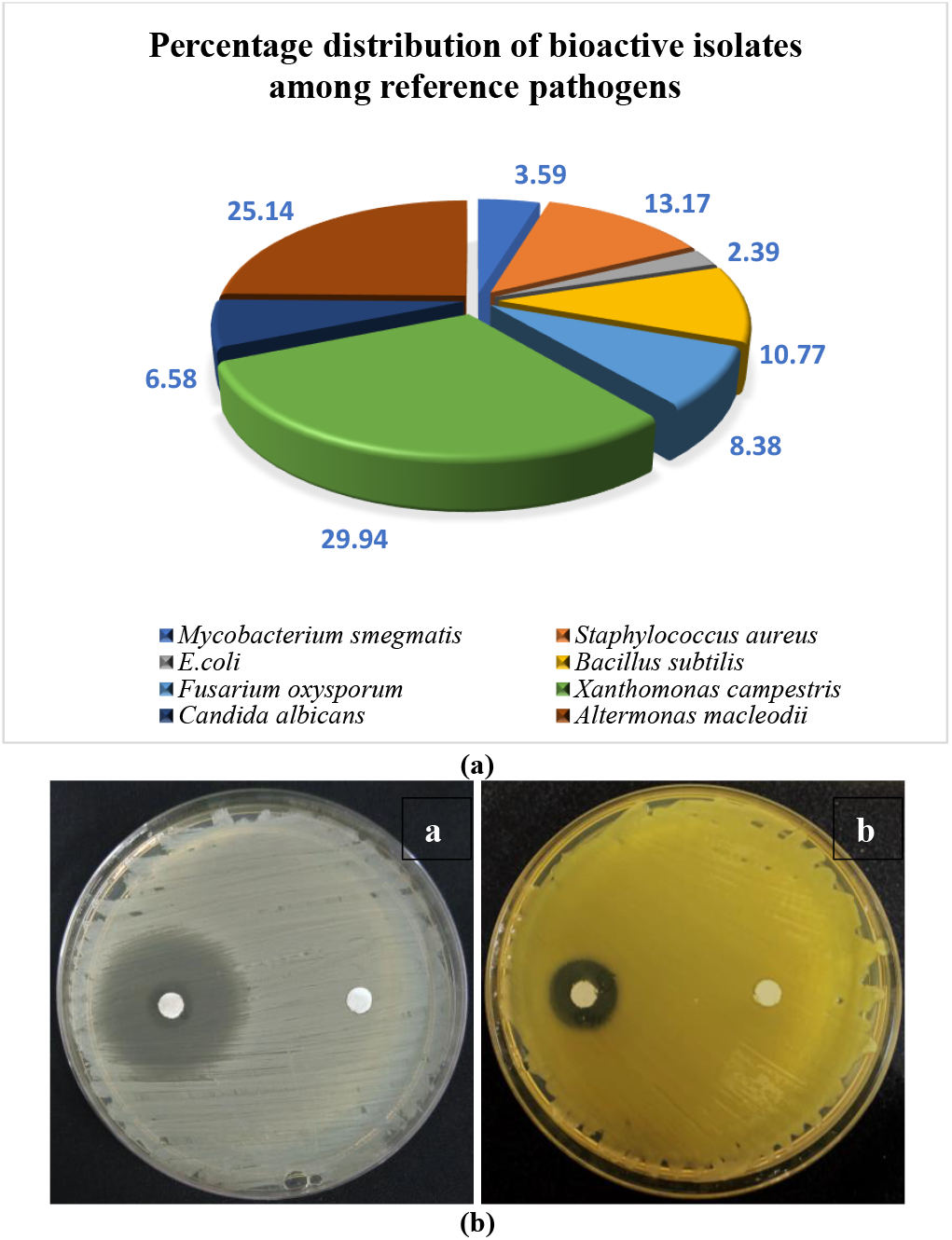
(a) Pie graph depicting distribution of isolates exhibiting bioactivity against various pathogens. (b) Bioactivity of *B. amyloliquefaciens* 5NPA-1 against (a) *S. aureus* MCC 2043 and (b) *X. campestris* NCIM 5028

**Table IV.**
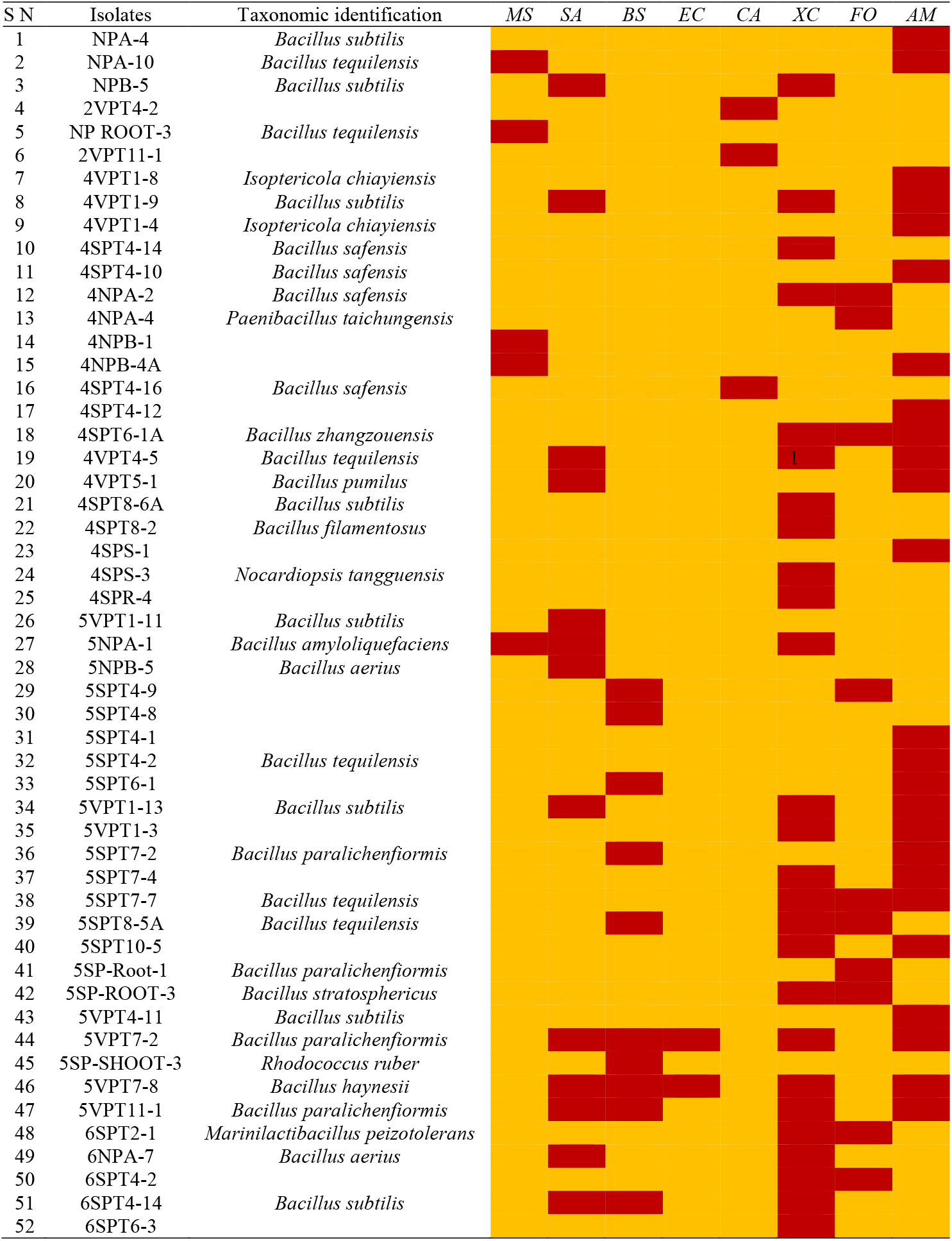
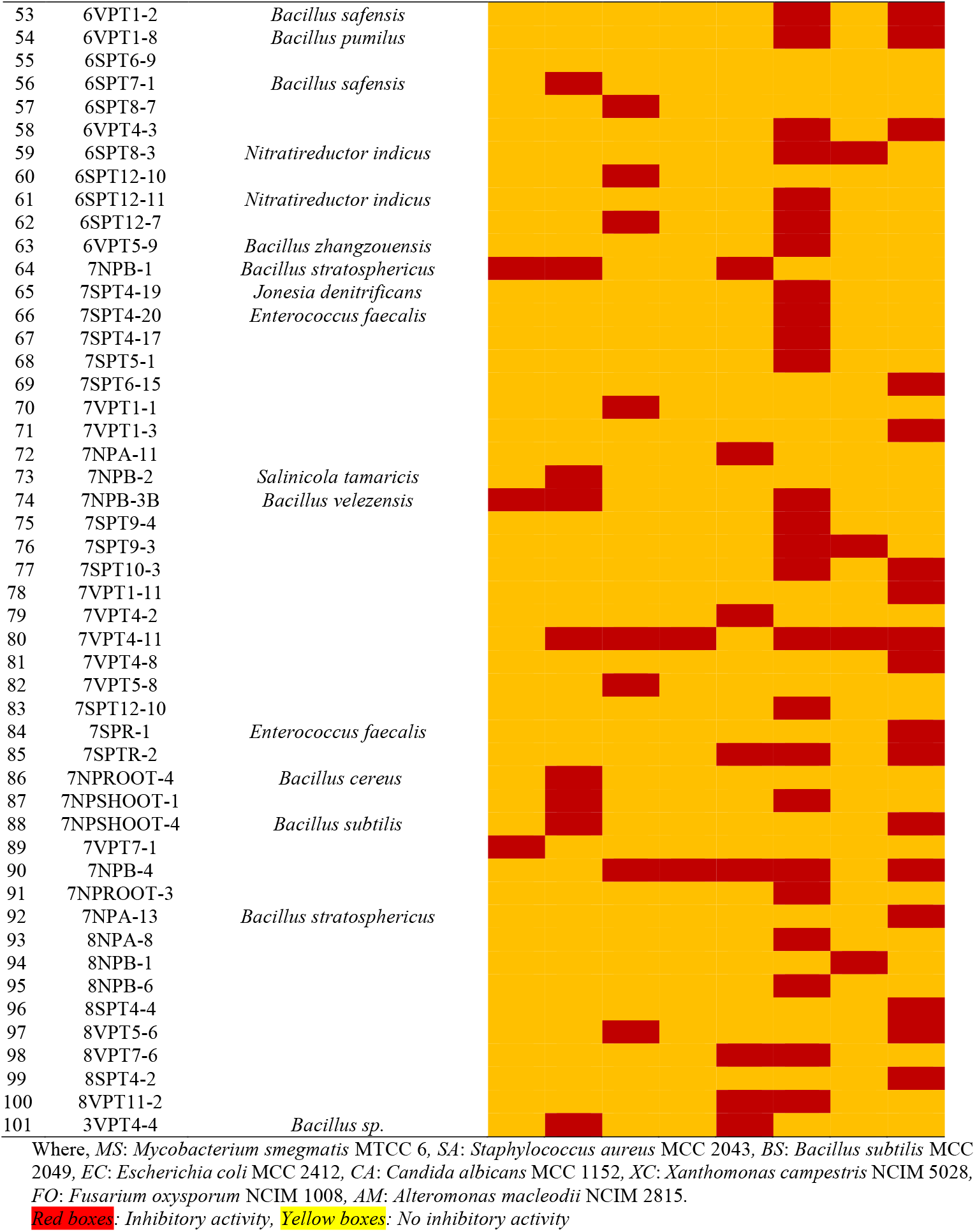
Results of primary screening of endophytic isolates against microbial pathogens.

### Preparation of crude extracts and

Crude extract of bioactive strains were prepared through solvent-solvent extraction method. Bioactivity of isolates active against *S. aureus* MCC 2043 and *X. campestris* NCIM 5028 was confirmed by disk diffusion assay (**Fig. 3b**), according to the guidelines of CLSI. The isolate *Bacillus amyloliquefaciens* 5NPA-1 exhibited a prominent zone of inhibition of 29 mm and 14 mm against pathogens *S. aureus* MCC 2043 and *X. campestris* NCIM 5028 respectively, serving as a potential isolate for isolation of bioactive compounds which are beneficial to plants as well as humans.

### LC-MS/MS based molecular networking

Molecular networking of crude extract of *B. amyloliquefaciens* 5NPA-1 using the GNPS platform was found to consist of 24 nodes grouped into 4 clusters. Largest cluster had 5 nodes, which was annotated as surfactin by automatic dereplication using MS/MS spectral libraries available at GNPS (**Fig. 4**). The network resulted into identification of 3 types of surfactin with respect to variations in length of fatty acid chain. Further, no specific networks denoting other class compounds were observed, supporting the idea that lipopeptides in *B. amyloliquefaciens* 5NPA-1 could be responsible for the bioactivity.

**Fig. 4.**
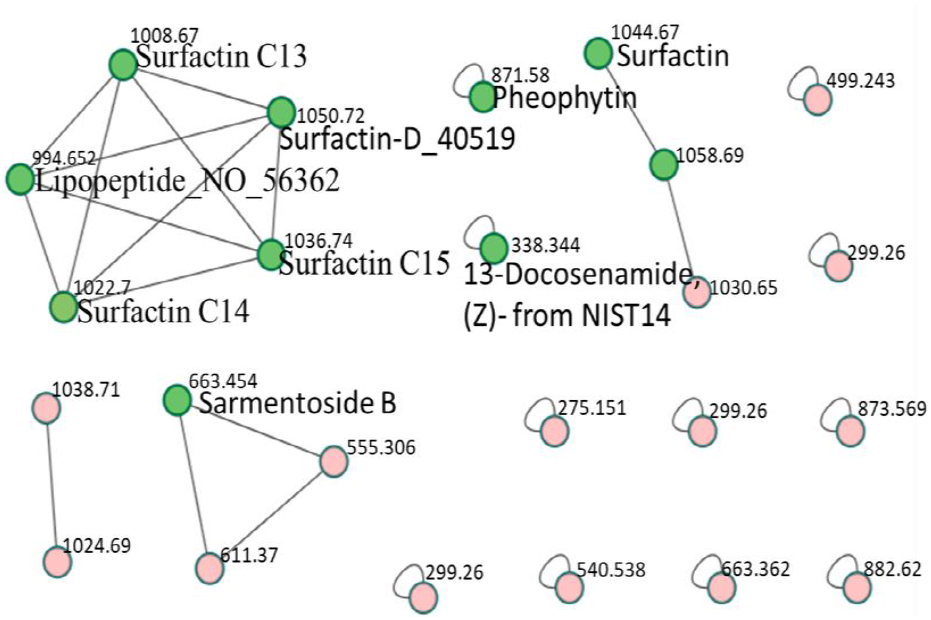
Molecular network of *B. amyloliquefaciens* 5NPA-1. Annotated molecular network (green) by GNPS for crude extract identifies surfactin networks. Molecular weights beside nodes indicate mass of parent ions. Unidentified nodes (pink) were not automatically detected.

### TLC Bioautography

In order to identify bioactive metabolites, TLC plate was developed to obtain 7–8 bands as observed under short UV radiations (254 nm wavelength). Developed TLC plate when overlaid with *S. aureus* MCC 2043 suspension on 1% agar, displayed clear zone with no cell growth against pinkish background at R_f_ between 0.12 to 0.41 (**Fig. 5**).

**Fig. 5.**
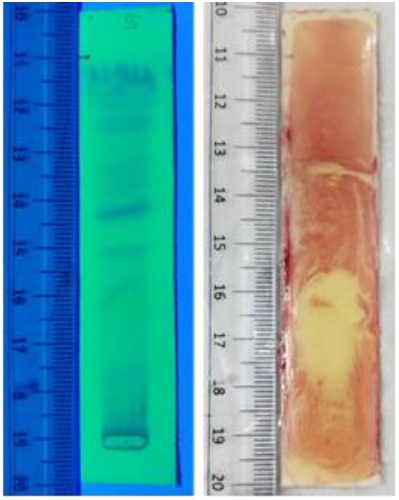
TLC bioautography plate of crude extract of *B. amyloliquefaciens* 5NPA-1. Pink area represents cell growth and clear zone area depicts presence of bioactive compound at that region

### Isolation and identification of bioactive molecules

The active middle band (5-PTLC-2) obtained from TLC-bioautography was scrapped in order to acquire ^1^H-NMR spectrum to identify the chemical class of the compounds. ^1^H-NMR spectrum of PTLC-2 exhibited signals for a long aliphatic alkyl chain at *δ*_H_ 1.29, CH_3_ groups at *δ*_H_ 0.85–1.00, an oxy-methine at *δ*_H_ 5.31, seven *α*-H at *δ*_H_ 4.05-4.80, indicating lipopeptide nature of compounds present in the fraction PTLC2 (**Fig. 6**). HPLC-DAD chromatogram of PTLC-2 revealed three peaks suggesting it to be a mixture of three compounds (**Fig. S3**). Hence, for their targeted separation, the crude ethyl acetate (EtOAc) extract was subjected to MPLC followed by HPLC (**Fig. S4**) resulting three compounds **1**, **2**, and **3** (**Fig. 7a**). The MALDI-TOF-MS spectra of **1**, **2**, and **3** exhibited intense sodium adduct [M+Na]^+^ ion peaks at *m/z* 1031.2147, 1045.0100 and 1058.8456, respectively. Comparison of the MALDI-TOF-MS data of **1**, **2**, and **3** with those available in literature revealed that the purified compounds belonged to surfactin class with different number of CH_2_ units in fatty acid chain and subsequently identified as C_13_-surfactin (**1**), C_14_-surfactin (**2**), and C15-surfactin (**3**) [27] (**Fig. S5,S6,S7**). To determine the sequence of the amino acids, the sodiated ions of **1**, **2**, and **3** were separately subjected to tandem MS experiment giving rise to spectra with diagnostic ion series containing C- and N-terminus. The MS/MS spectra of compounds **1**, **2**, and **3** displayed a major ion sequence of fragments *m/z* 707.4, 594.3, 481.2, 382.1, 267.1 corresponding to loss of (fatty acid)-Glu-Leu-Leu-Val-Asp-Leu-Leu from C-terminal region. Similarly, another major fragment ion series of **1** [*m/z* 945.5, 832.5, 717.4, 618.4], **2** [*m/z* 931.5, 818.5, 702.4, 604.3], and **3** [*m/z* 917.5, 804.4, 688.4, 590.3] confirmed the presence of amino acid residues sequence as Leu-Leu-Asp-Val-Leu-Leu-Glu-(fatty acid) from the N-terminal region. Taken together, the connection of the two series suggested that the surfactins **1**, **2**, and **3** contained the same heptapeptide sequence (fatty acid)-Glu-Leu-Leu-Val-Asp-Leu-Leu (**Fig. S8,S9,S10**). ^13^C NMR chemical shifts allowed to differentiate branching of the hydroxy fatty acid side chain among the *normal (*δ*_C_* 13.8, 22.0, 31.2), *iso* (*δ*_C_ 22.4, 22.4, 27.3, 38.4), and the *anteiso* (*δ*_C_ 11.1, and 19.0) chain types [28]. Based on this approach, the *β*-hydroxyl fatty acid chains were found to be mixtures of *iso*-C_10_H_21_ and *anteiso*-C_10_H_21_ in **1**; *n*-C_11_H_23_, *iso*-C_11_H_23_, and *anteiso*-C_11_H_23_ in **2**; and, *iso*-C_12_H_25_ and *anteiso*-C_12_H_25_ in **3**, respectively (**Fig. 7b**). On the basis of the previous studies, absolute configurations of amino acid units from N- to C-terminal of **1**, **2**, and **3** were assumed to be L-, L-, D-, L-, L-, D- and L-, respectively, and the C-3 configuration of fatty acid was assumed to be as *R* [29].

**Fig. 6.**
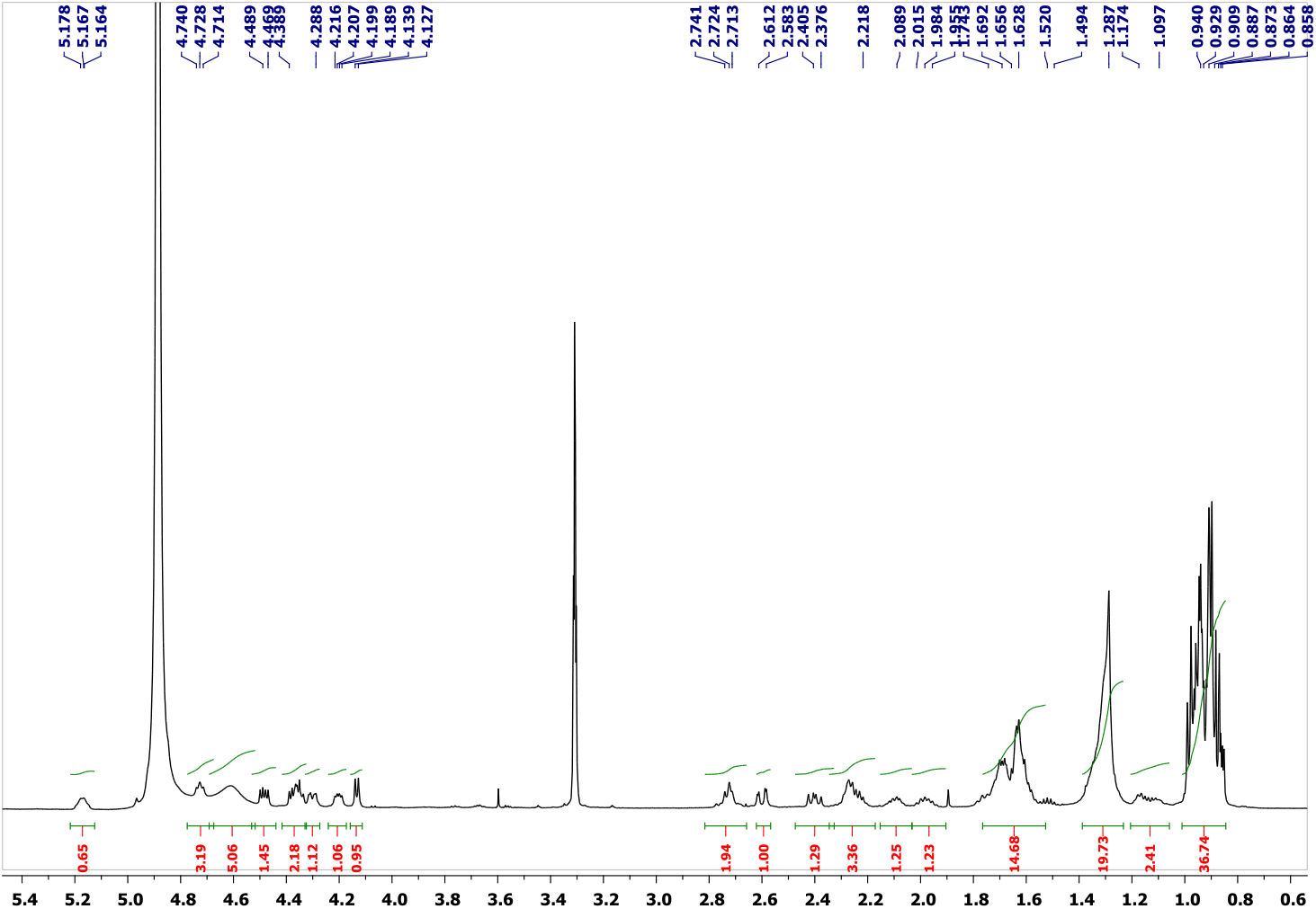
^1^H-NMR spectrum of 5-PTLC2 fraction obtained from preparative TLC of crude extract of *B. amyloliquefaciens* 5NPA-1

**Fig. 7.**
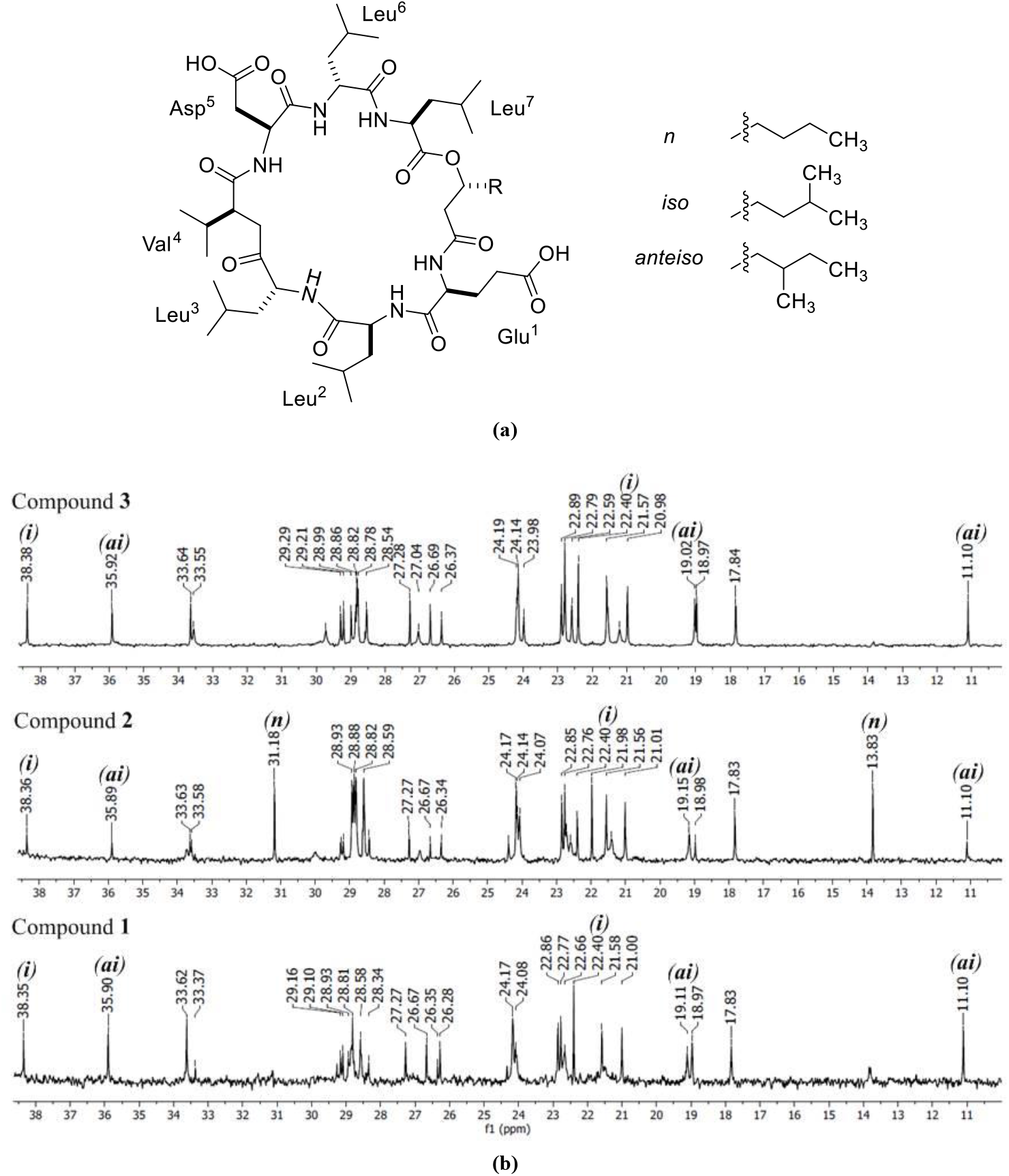
(**a**) Structures of compounds **1** (R = mixture of *iso*-C_10_H_21_ and *anteiso*-C_10_H_21_ patterns), **2** (R = mixture of *n*-C_11_H_23_, *iso*-C_11_H_23_, and *anteiso*-C_11_H_23_ patterns), and **3** (R = mixture of *iso*-C_12_H_25_ and *anteiso*-C_12_H_25_ patterns). (**b**) Characteristic ^13^C NMR signals differentiating branching of the hydroxy fatty acid side chains in compounds 1–3; (**n**): normal (*δ*_C_ 13.8, 22.0, 31.2), (**i**): iso (*δ*_C_ 22.4, 22.4, 27.3, 38.4) and the (**ai**): anteiso (*δ*_C_ 11.1, and 19.0) chain type.

## DISCUSSION

Endophytes, plant associated symbionts, have emerged as an interesting source for natural products because of their diversity in bioactive secondary metabolites. Halophytes are reported to overcome their abiotic and biotic stress with the help of metabolites, regulators or enzymes released from endophytes [30]. In this study, diversity of bacterial endophytes from *S. brachiata* was assessed and their probable role in host-plant interaction including identification of metabolites produced by one of the endophyte *B. amyloliquefaciens*. All endophytes were screened for antimicrobial potential against human, plant, and marine pathogens. To our knowledge this is a first comprehensive report regarding diversity of bacterial endophytes from *Salicornia brachiata* from Gujarat coast of India and investigation of their antimicrobial potential.

Surface sterilized plant material with six different pretreatments and eight different previously reported media resulted into a total of 336 endophytes delivering a quantitative idea about the bacterial diversity within the halophyte. The results indicated a mixed composition of the endophyte communities comprising majorly firmicutes followed by actinobacteria and proteobacteria. Studies on endophytic bacteria from different parts of halophytes *Salicornia europaea, Arthrocnemum macrostachyum* etc. have been performed previously and predominance of aforementioned phyla was observed [31,32]. Out of 336 endophytes, 63 isolates identified in the study represented 13 genera with 29 different species. *Bacillus* as a dominant genus was observed with a diversity of around 15 different species. *Bacillus* sp. due to their better resilience is usually the dominant firmicute isolated from saline environments [30,33]. The statistical analysis showed a higher richness and evenness of species diversity among isolated endophytes. Amidst the endophytes obtained in our study from *S. brachiata*, genera *Bacillus, Isoptericola, Streptomyces, Salinicola, Rhodococcus* have already been reported from sister species of *Salicornia europaea* [32,34]. *Nocardiopsis, Jonesia, Nitratireductor, Paenibacillus, Micromonospora* are some genera from *S. brachiata* we report in our study.

It is well established that endophytes support plant ecological progression through production of various metabolites. Processes supported by these metabolites increases bioavailability of nutrients to host, tolerance against abiotic stress and strength to fight against biotic stress including pests and phytopathogens. Among the isolated strains in the study *Bacillus* sp. is predominantly reported to produce ACC deaminase enzymes to alleviate stress by ethylene, indole acetic acid and gibberellic acids promoting cell division and growth [30], phosphate solubilization enzymes, biological nitrogen fixation, siderophores and bioactive metabolites against phytopathogens [5,31,35]. Bioactive metabolites from *Bacillus* sp. include polyketides bacillomycin, fengycin, iturin, lichenysin, mycosubtilin, plipastatin, pumilacidin, and surfactin [36]. Hence, abundance of *Bacillus* endophytes can be validated due to its profuse chemical interactions with host plant. The strains *Salinicola sp*. and *Rhodococcus sp*. also display ACC deaminase activity important for plant growth promotion events in stress conditions [34]. Moreover, endophytic *Salinicola* sp. isolated from *Spartina maritima* was reported to be an excellent producer of siderophores and contain heavy metal tolerant genes thereby supporting the plant to alleviate the toxic effect of heavy metals [37]. The actinobacterial genera of *Nocardiopsis* and *Isoptericola* [38] thrive in the saline conditions. Genetic makeup of *Nocardiopsis* is filled with megaplasmid genes encoding antibiotic productions like apoptolidin, lipopeptide biosurfactants, thiopeptides, griseusin D etc., heavy metal resistance and stress response including osmoregulation benefitting their survival in halophilic environment [39]. It can be said that the ecological stress within host plant stimulated production of such bioactive metabolites in *Nocardiopsis. Jonesia denitrificans* as the name suggests is reported to perform denitrification [40]. Various strains of *Streptomyces* are reported to exhibit phosphate solubilization property, ammonia production, enzymes production for breakdown of organic matter, PKS and NRPS gene clusters for production of bioactive compounds etc. which contributes to plant health either directly or indirectly [41]. These reports reflect direct plant-microbe interactions of the isolates obtained in study, further supporting their endophytic origin from halophyte.

In-vitro screening of the isolates for bioactivity revealed one third of the population to be bioactive against one or more reference pathogen. This aligns with the work of Verma *et al*. who reported 60% of the endophytic actinobacteria isolates obtained from *Azadirachta indica* showed wide-spectrum antagonistic potential [42]. Given their metabolite productions, a large number of isolates exhibited inhibition of growth of plant pathogens *X. campestris* NCIM 5028, *F. oxysporum* NCIM 1008 and marine bacterial pathogen *A. macleodii* NCIM 2815. Inhibition of phytopathogens at such enormous amount indicates role of endophytes in defense mechanisms of host plant. Some of the isolates were found to inhibit *M. smegmatis* MTCC 6, *S. aureus* MCC 2043, etc., suggesting that the antimicrobial activity exhibited by plant *S. brachiata* [13] can be attributed to the bioactive metabolites secreted by inhabiting endophytes. The genus *Bacillus* was dominant in displaying activity against all indicator pathogens pertaining to its siderophore and bioactive lipopeptide production potential. Production of siderophores was reported from halotolerant *Bacillus* isolated from wheat seedlings, further it improve soil fertility increasing plant productivity in agriculture and also remediates toxic metals from human body [43]. *Bacillus* sp. have been isolated as endophytes from ginger, turmeric etc. and shows enormous antifungal properties and antibacterial properties due to the presence of cyclic lipopeptides [44].

In the present work, along with endophytic biodiversity, an emphasis was also laid on isolate displaying activity against *S. aureus* MCC 2043, a common nosocomial pathogen and *X. campestris* NCIM 5028, a meticulous plant pathogen. Crude extract of isolate *B. amyloliquefaciens* 5NPA-1 (MT459305) was found to be most potent with zone of inhibition of 29 mm and 14 mm against *S. aureus* MCC 2043 and *X. campestris* NCIM 5028 respectively. Molecular networking using LC-MS/MS data of crude extract gave an idea about the presence of secondary metabolites encrypted in the strain *B. amyloliquefaciens* 5NPA-1 indicating surfactin type compounds. High potency of crude extract served as a driving force for purification of bioactive compounds to identify its potential as strong antimicrobial agents. Following that paradigm, purification and characterization of active compounds from *B. amyloliquefaciens* 5NPA-1 was also performed leading to compounds **1**–**3** belonging to lipopeptides class. Lipopeptides form an important class of metabolites from endophytic bacteria, wherein serving as antibiotic and inducing plant systemic resistance. *B. amyloliquefaciens* was recognized as a higher lipopeptide producer when isolated from different plants including *Phaseolus vulgaris, Oryza sativa, Ophiopogon japonicus, Musa acuminata*, marine plants etc., meanwhile also secreting plant growth promoters, phyto-hormones, siderophores, antifungal, anticancer and antimicrobial agents [36]. Diversity of endophytes within plant structures is proportional to various benefits of plant-microbe interactions. Such interactions are of ecological importance as they improve adaptation capabilities of either species and improve soil fertility and texture. An understanding of the chemical ecology of plants-microorganisms should enable the development of new crop improvement strategies, the conservation of indigenous varieties, and definitely a source of interesting pharmaceutical compounds.

To conclude, present study was the first attempt where endophytic bacterial community residing in stress-tolerant halophyte *S. brachiata* was studied and examined for the production of antimicrobial compounds against pathogens of various niches. Through identification of 20 % isolates, it was revealed that the plant harbors a rich bacterial biodiversity accounting for 13 genera and 29 species with *Bacillus* being dominant and distinct actinobacteria exhibiting different morphology, producing pigments, metabolites and polysaccharides which benefits the plant. Metabolites from species inhabiting the plant have history in supporting the host plant through various chemical interactions. It was also deciphered that surfactin class molecules produced by endophytic strain *B. amyloliquefaciens* 5NPA-1 possess high biocontrol properties against nosocomial pathogen and bacterial plant pathogen. *Bacillus amyloliquefaciens* being an environmentally stable and fast replicating bacteria serves as an ideal source for extraction of plethora of metabolites. Such enormous antimicrobial potencies displayed by several endophytes from *S. brachiata* indicate their role in plant defense system, and serve as an example of plant microbe interaction. This diverse population can be further explored for novel metabolites given that demand for novel bioactive agents is everlasting and it may help us with better understanding the chemical ecology of an ecosystem.

## Data Availability Statement

The datasets generated for this study are available on request to corresponding author.

## Author Contributions

Sanju Singh, Vishal Ghadge, Pankaj Kumar, and Pramod B. Shinde designed and planned the research. Sanju Singh, Vishal Ghadge, and Pankaj Kumar isolated and identified the bacterial strains. Sanju Singh and Doniya Elze Mathew performed bioactivity. Pankaj Kumar performed molecular networking analysis. Sanju Singh, Asmita Dhimmar, Pankaj Kumar, Harshal Sahastrabudhe, and Yedukondalu Nalli isolated and identified bioactive secondary metabolites. All authors analyzed and interpreted the results and commented on the manuscript prepared by Sanju Singh and Pramod B. Shinde.

## Acknowledgments

Sanju Singh, Doniya Elze Mathew, Asmita Dhimmar acknowledges CSIR-JRF fellowship from Council of Scientific and Industrial Research (CSIR), Pankaj Kumar acknowledges DBT-JRF fellowship from Department of Biotechnology (DBT) and Harshal Sahastrabudhe acknowledges GATE-JRF fellowship from Council of Scientific and Industrial Research (CSIR). Yedukondalu Nalli acknowledges Scientific and Engineering Research Board (SERB) for providing National Postdoctoral Fellowship (N-PDF).

## Supplementary Material

A schematic diagrams of isolation of endophytes and compounds, HPLC chromatograms, fragmentation pattern, and MS- and MS/MS-spectra of compounds **1**–**3** is available.

